# Improving cell-type identification with generative adversarial networks-enhanced augmentation-free single-cell RNA-Seq contrastive learning

**DOI:** 10.64898/2025.11.29.691320

**Authors:** Ibrahim Alsaggaf, Daniel Buchan, Cen Wan

## Abstract

Cell-type identification plays a fundamental role in single-cell RNA-Seq analytics. Thanks to the recent success of the contrastive learning paradigm, the accuracy of automatic cell-type identification has also been improved. In this work, we propose a novel contrastive learning-based cell-type identification method, namely GAN-RCL, which conducts hard positive sample selection on real and conditional generative adversarial networks-created synthetic instances to improve the performance of augmentation-free contrastive learning. Experimental results confirm that GAN-RCL successfully outperforms other recently-proposed contrastive learning-based methods and shows the state-of-the-art predictive performance on multiple single-cell RNA-Seq cell-type identification tasks.

## 1 Introduction

Single-cell RNA sequencing (scRNA-Seq) is an important experimental methodology in contemporary transcriptomics that allows researchers to sequence the transcriptome of single cells [1] to obtain an informative transcriptomic landscape. Accurate cell-type identification plays a fundamental role in scRNA-Seq analytics. However, the well-known high sparsity and high dimensionality characteristics of scRNA-Seq expression profiles are still considered as the major challenges for the conventional machine learning-based scRNA-Seq analytics methods. Thanks to the recent development of the self-supervised learning paradigm, many contrastive learning-based scRNA-Seq analytics methods were proposed and obtained state-of-the-art performance on the variety of scRNA-Seq analytics tasks. For example, Ciortan and Defrance (2021) [3] first proposed a random genes masking-based contrastive learning method for scRNA-Seq clustering tasks, which were also studied by Wan et al. (2022) [4] to develop a more sophisticated neighborhood contrastive clustering method. Xu et al. (2022) [5] proposed a contrastive learning-based method that can integrate multisource single-cell transcriptome data whilst remove batch effects. Yang et al. (2022) also developed a contrastive learning method for integrating and projecting multimodal single-cell atlases into a type of hypersphere feature space. More recently, Alsaggaf et al. (2024) [6] first proposed a Gaussian noise augmentation-based cell-type identification method that successfully outperformed other conventional machine learning-based methods, such as the well-known scPred [7] and ACTINN [8]. In 2025, Alsaggaf et al. [9] further proposed an augmentation-free contrastive learning method that obtained the state-of-the-art performance on the variety of cell-type identification tasks.

Contrastive learning aims to learn a type of discriminative distributions where similar instances are pulled closer and dissimilar instances are pushed away. The conventional self-supervised contrastive learning methods, such as SimCLR [10], first create two views for each instance (a.k.a. an anchor) by using different data augmentation strategies. For each anchor, the corresponding two views are treated as a positive sample and all other instances are treated as a negative sample. Then SimCLR optimises the network parameters in order to reduce the distance between those two instances in the positive sample, whilst enlarges the distance between the positive sample and the negative sample. The self-supervised learning paradigm was further extended to the supervised contrastive learning paradigm [11], where the definition of positive and negative samples relies on the original labels of instances. For each anchor, the positive sample consists of those instances whose labels are the same as the anchor’s. *Vice versa*, the negative sample consists of those instances bearing different labels to the anchor. This supervised contrastive learning paradigm successfully demonstrated better predictive performance than the conventional self-supervised contrastive learning paradigm on the variety of tasks.

Data view creation and selection play crucial roles in the performance of contrastive learning, since networks’ optimization gradients are derived by computing the similarity between views in positive and negative samples. For example, in terms of image-based contrastive learning tasks, conventional data augmentation approaches, such as colour distortions and image cropping [10], are commonly used to create views. However, different data augmentation approaches usually lead to stronger or weaker views. It has been shown by [12, 13] that an appropriate combination of strong and weak positive views is required to improve the performance of contrastive learning with respect to downstream tasks. In terms of views in the negative sample, due to its larger quantity, compared with positive views, the view selection strategies were also further studied with an aim to create a better negative sample including hard data views. For example, the so-called hard negatives are defined as those views belong to different classes to the anchor but have high similarities with the anchor. It has been recently confirmed by different works [14, 15] that hard negatives selection strategies can successfully improve the performance of contrastive learning.

Generative Adversarial Networks (GAN) are a type of generative model that can accurately capture high-dimensional distribution by adopting a type of zero-sum game paradigm. A classic GAN architecture is composed of two neural networks, i.e. a generator and a discriminator. The former generates fake samples by learning target distributions, whilst the latter tries to discriminate between fake and real training samples. The parameters of GAN are optimised until both the generator and the discriminator reach the Nash equilibrium where the fake and real samples are highly similar and cannot be well classified. Due to the superb capacity for learning high-dimensional distributions, GAN has already been used as a data augmentation approach. For example, Wan and Jones [16] used GAN to generate fake samples of high-dimensional protein biophysical properties. The fake samples successfully improve the accuracy of *Drosophila* protein function predictions by enriching the training dataset, which leads to improved support vector machine decision boundaries. Marouf, et al. [17] also used conditional GAN to generate synthetic single-cell RNA-Seq samples that improved the clustering quality of rare cells and improved the performance of marker gene detection. In this work, we propose a novel Generative Adversarial Networks-enhanced scRNA-Seq Contrastive Learning method, namely GAN-RCL. Computational experiments confirm that GAN-RCL successfully improves the performance of the state-of-the-art AF-RCL cell-type identification method [9], which was recently developed by us. The paper is organised as follows. The Background section briefly introduces contrastive learning, hard instance selection and conditional generative adversarial networks, along with a table showing the definitions of all notations used. The Methods section introduces the proposed GAN-RCL algorithm, followed by the Computational Experiments section, where the Experimental Results subsection reports the performance of GAN-RCL and compares it with other cell-type identification methods. Finally, the Conclusion and Future Research Directions section summaries the contributions of this paper and reveals future research directions.

## 2 Background

### 2.1 Contrastive Learning Paradigm

Contrastive learning aims to learn a hypersphere feature space where similar instances are pulled closer and dissimilar instances are pushed further away. Conventional contrastive learning follows a form of selfsupervised learning paradigm [10], i.e. the learning gradients of networks optimisation are generated by comparing training instances’ views without exploiting their corresponding class label information. In 2020, Khosla et al. [11] extended the conventional self-supervised contrastive learning paradigm to the supervised learning case, this seeks to pull instances bearing the same class label closer whilst push those instances bearing different class labels further away. This extension successfully improves the performance of contrastive learning methods in general.

**Table 1.**
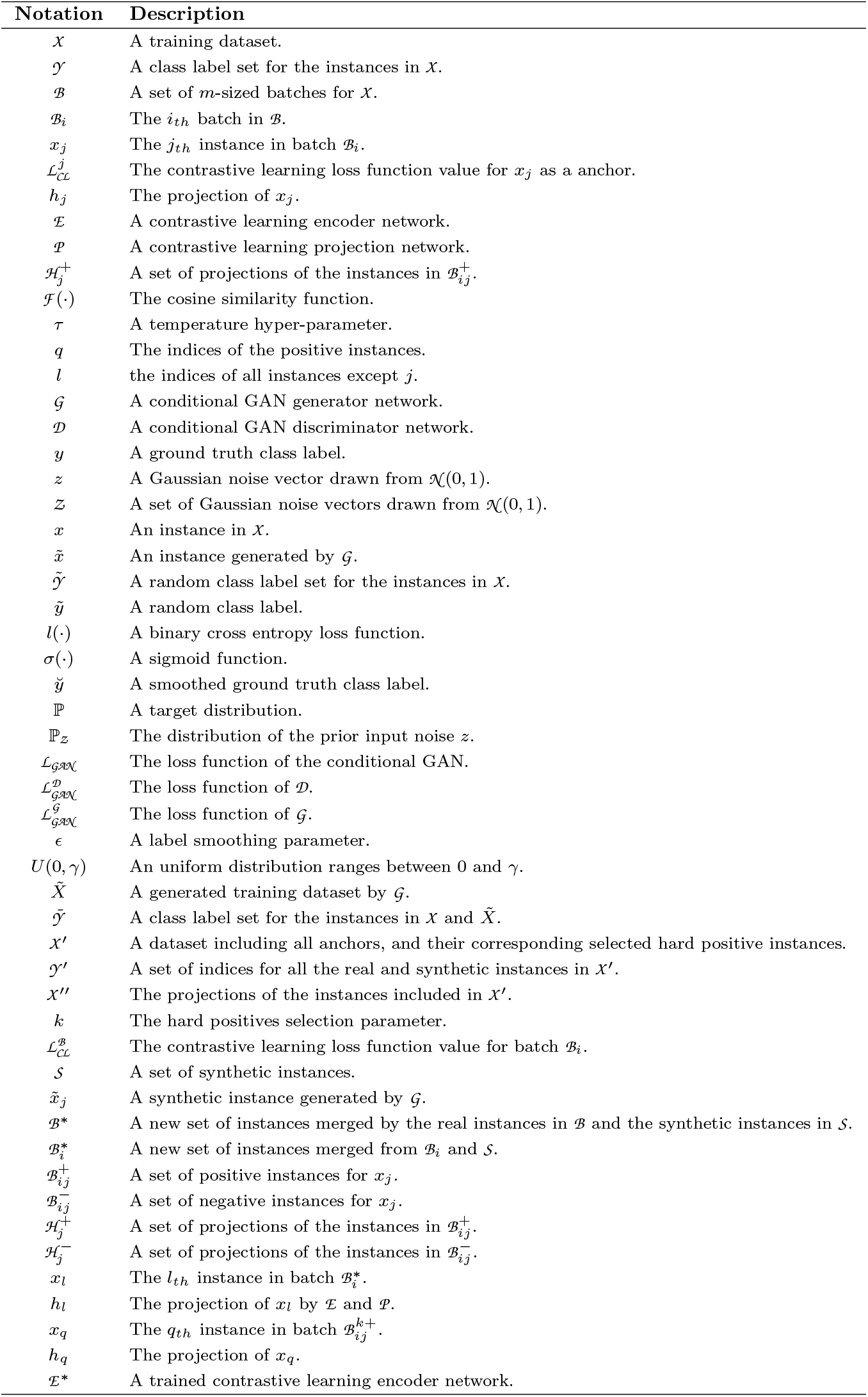
The list of notations used in this paper.

Both self-supervised and supervised contrastive learning methods have already been used for the scRNA-Seq cell-type identification tasks. In 2024, Alsaggaf et al. [6] proposed four new scRNA-Seq contrastive learning-based cell-type identification methods, i.e. Self-GsRCL, Self-RGMRCL, Sup-GsRCL and Sup-RGMRCL. Self-GsRCL and Self-RGMRCL obtained the state-of-the-art predictive accuracy on difficult and easy cell-type identification tasks, respectively. In 2025, Alsaggaf et al. [9] further proposed an augmentation-free contrastive learning method (i.e. AF-RCL), which successfully further improved the predictive accuracy of cell-type identification tasks in general. AF-RCL follows the conventional supervised contrastive learning paradigm, but it directly exploits original training instances during training without creating any views. Hence, AF-RCL directly pulls closer those cells having the same cell-types and pushes further away those cells belonging to different cell-types. In addition to the new augmentation-free setting, AF-RCL also exploited a new loss function, as shown in Equation 1. *h*_*j*_ denotes the latent features of the anchor *x*_*j*_ (i.e. *h*_*j*_ = 𝒫 (ℰ (*x*_*j*_))) and 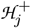 denotes the transformed positive instance set. ℱ (·) denotes the cosine similarity and *τ* is a temperature hyperparameter. *q* indicates the indices of the positive instances and *l* indicates the indices of all instances except *j*.

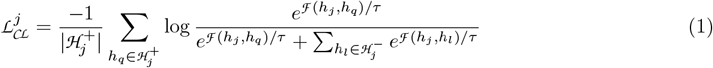

### 2.2 Hard Instance Selection for Contrastive Learning

As the gradients of contrastive learning networks optimization are derived by considering the similarity between views of training instances, the performance of contrastive learning can be improved by adopting stronger data augmentation view creation approaches or by selecting more informative views. In terms of the positive instances with respect to one anchor instance, the most commonly used data augmentation approaches for image data are Gaussian blurring, rotation and color-jittering [10]. However, excessively strong data augmentation approaches would lead to advert impact on downstream tasks, due to overfitting issues and information loss. Therefore, many methods were recently proposed to integrate views created by strong and weak augmentation approaches [12, 13, 18]. In terms of the negative instances with respect to one anchor instance, it is common to adopt instance selection strategy to create a so-called hard negative set, since the numbers of negative instances are usually much larger than positive instances. For example, Robinson, et al. (2021) [14] proposed a group of hard negatives selection methods for the conventional self-supervised contrastive learning, whilst Jiang, et al. (2024) [15] also proposed a class-conditioned negatives selection method for the supervised contrastive learning paradigm. As discussed in our previous work [9], supervised augmentation-free contrastive learning method (i.e. AF-RCL) outperformed other augmentation-based contrastive learning methods for cell-type identification tasks. Therefore, in this work, we focus on improving the quality of positive instances with respective to anchors without adopting any augmentation-based view creation approaches.

### 2.3 Conditional Generative Adversarial Networks

Conditional GAN (cGAN) is a variant of GAN where a pair of generator 𝒢 and discriminator 𝒟 networks are trained conditioned on auxiliary information, such as class labels [19]. Equation 2 shows the loss function of the class label-based conditional GAN used in this work, where ℙ is the target data distribution and ℙ_*z*_ is the distribution where random noises can be drawn. 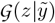 denotes an instance created by the generator using a prior noise *z* and a random class label value 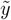. Note that, in order to improve the cGAN training, we use a random class label value 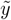 instead of the ground truth class label *y*, which is used in the conventional cGAN [19]. In addition, due to the training stability concerns [20], in this work we use Equation 3 as the loss function along with the label smoothing approach to improve models’ generalisability [21–23]. The loss function exploits the well-known binary cross entropy loss function *l*(·), whilst *σ*(*·*) denotes a sigmoid function. Suppose that *y* is the ground truth class label, whilst 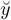 is the smoothed ground truth class label, the discriminator is trained to minimise Equation 3, where 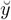 is close to 1 in the first term and 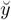 is close to 0 in the second term. To enforce the adversarial training, the generator maximises the second term in Equation 3 by minimising Equation 4 where we change 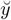 from 0 to 1 to fool the discriminator. In terms of the label smoothing approach, we simplify the mechanism to fit the binary case by letting *ϵ* be a smoothing parameter that is randomly drawn from an uniform distribution *U* (0, *γ*), where 0 *< γ <* 0.5. Let 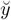 be the smoothed label of the hard ground truth *y*. As shown in Equation 5, we set 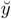 to a number close to 1 for real samples and to a number close to 0 for synthetic samples. Finally, analogous to the work in [16], we adopt Classifier Two-Sample Tests (CTST) [24] for model selection, i.e. the optimal generator is selected if it leads to a leave-one-out cross validation accuracy close to the chance-level, i.e. 0.500, denoting that the distribution of the target data is successfully learned by the generator network.

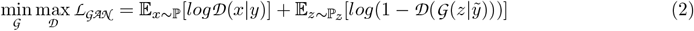

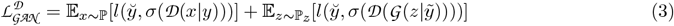

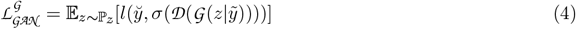

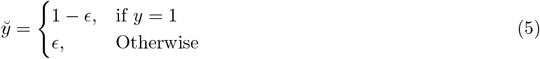

## 3 Proposed Methods

The proposed method, GAN-enhanced single-cell RNA-Seq Contrastive Learning (GAN-RCL), exploits synthetic samples generated using conditional GANs to enlarge the training sample space. With this method, more informative instances can be selected for conducting contrastive learning. Figure 1 shows an overview of GAN-RCL, which includes three components, i.e. a single-cell RNA-Seq-based generative adversarial networks (scGAN) training stage (Figure 1.A), a hard positive scRNA-Seq sample selection stage (Figure 1.B), and a scRNA-Seq contrastive learning stage (Figure 1.C). In the scGAN training stage, a conditional GAN is trained by using training scRNA-Seq instances and the corresponding class labels to generate synthetic scRNA-Seq instances (e.g.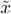) bearing different cell-type labels. In the hard positive scRNA-Seq sample selection stage, the synthetic instances are merged with the real scRNA-Seq instances (e.g. *x*) to create a larger combined sample set ℬ^∗^. Only the most informative scRNA-Seq instances are selected to be kept in the scRNA-Seq contrastive learning stage, which follows the same procedure of the previous proposed AF-RCL learning procedure.

**Fig. 1.**
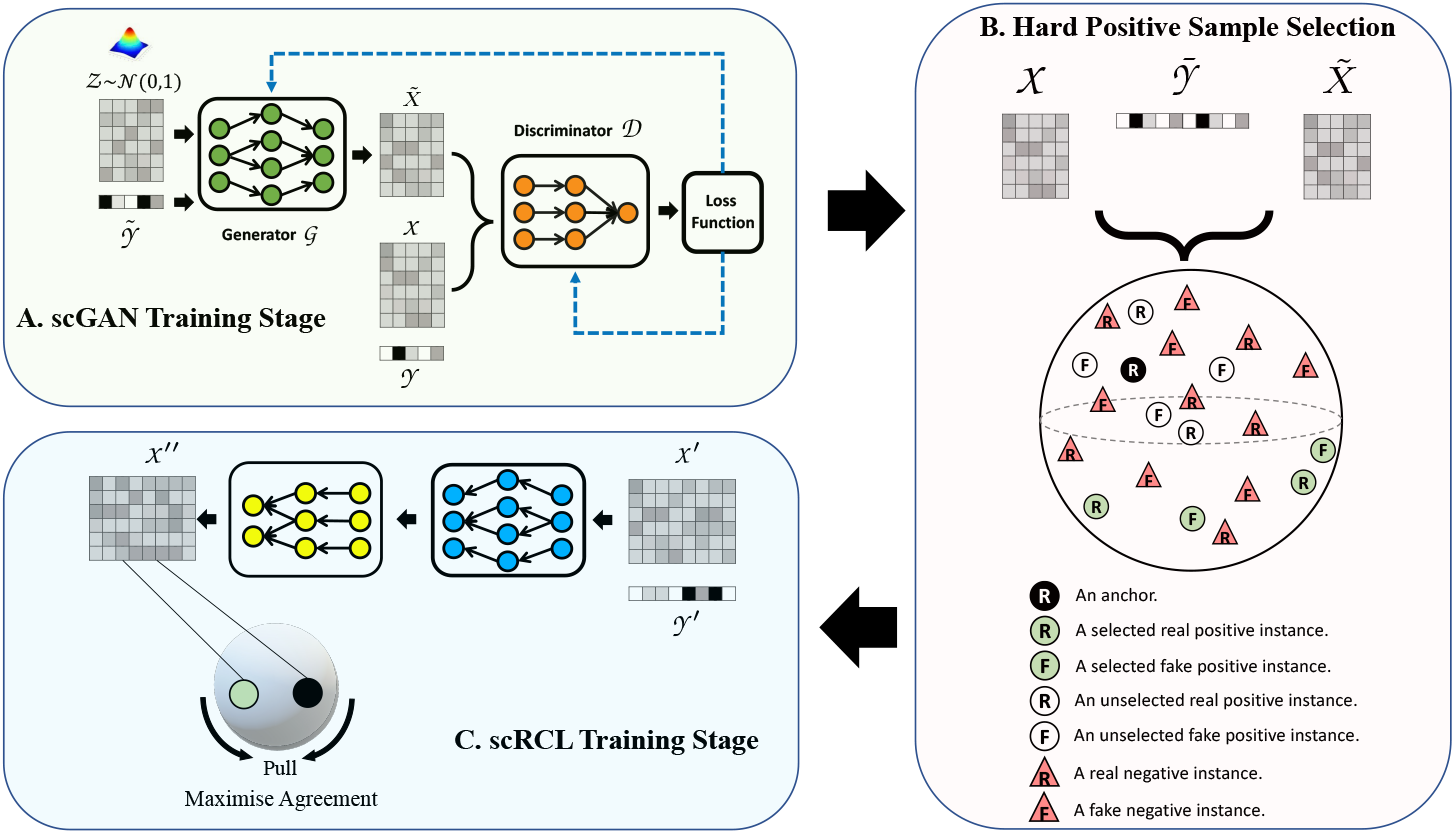
The flow-chart for GAN-RCL. The GAN-RCL method consists of three stages. (A) The first stage is using the original real scRNA-Seq expression profiles 𝒳 and their cell-types (i.e. class labels) 𝒴 to train a conditional GAN, namely scGAN. The trained scGAN is then used to generate synthetic scRNA-Seq expression profiles 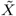. (B) The second stage is selecting hard positive instances. Both the real instances in 𝒳 and the synthetic instances in 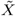 are merged as a new set, along with a new class label set 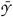 consisting of 𝒴 and its duplication. The hard positive instance selection process returns indices 𝒴′ of selected optimal hard positives w.r.t. each real instance as an anchor and all the selected real and synthetic instances as a set 𝒳′. (C) The third stage is conducing supervised augmentation-free contrastive learning, namely scRCL, by using 𝒳′ and 𝒴′. 𝒳″ denotes the transformed *𝒳*′ by using ℰ and 𝒫. The loss function 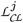 is used to pull similar instances closer, whereas push dissimilar instances away.

### Algorithm 1

Generative Adversarial Networks-enhanced Augmentation-free Single-cell RNA-seq Contrastive Learning (GAN-RCL)

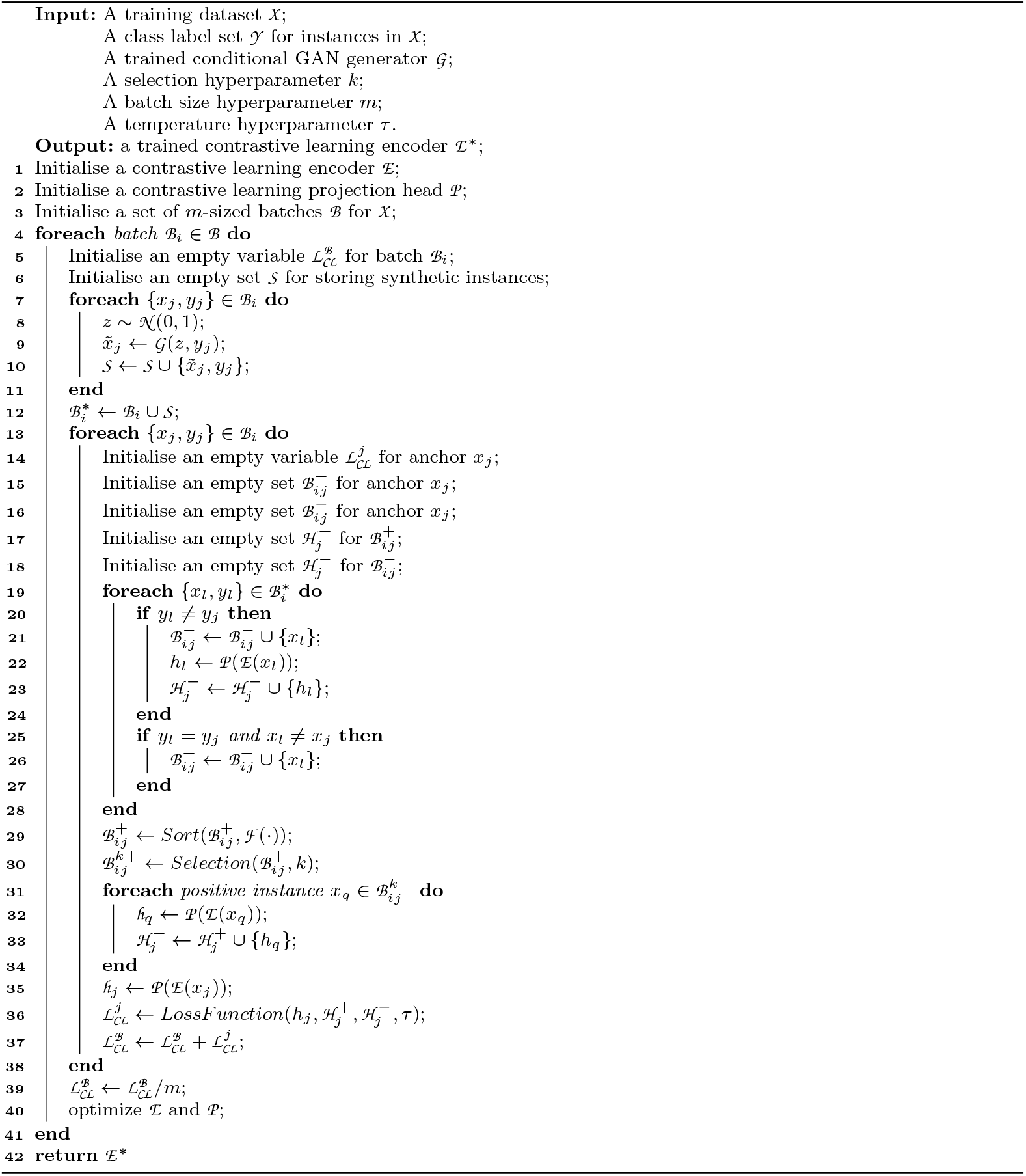

Algorithm 1 shows the pseudocode of the positive scRNA-Seq instance selection and contrastive learning processes. Given a training sample set 𝒳 with its corresponding label set 𝒴 and a corresponding scGAN that is already trained, Algorithm 1 returns a trained contrastive learning encoder ℰ^∗^ conditioned on three input hyperparameters *k, m* and *τ*. From line 1 to line 3, Algorithm 1 first initialises a pair of untrained contrastive learning encoder ℰ and projection head 𝒫, and a set of batches ℬ that individually consists of *m* instances. Then between lines 4 and 41, Algorithm 1 processes each batch ℬ_*i*_ to optimize the encoder ℰ and the projection head 𝒫. In terms of each batch, in lines 5 and 6, two empty variables 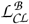 and 𝒮 are created to store the loss function value and the set of synthetic instances respectively. Between lines 7 and 11, for each real instance *x*_*j*_, a Gaussian noise is drawn and used as a input along with the label value *y*_*j*_ to generate a synthetic instance 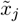 by using the trained scGAN 𝒢. The elements of 𝒮 is also updated by merging with 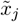. After processing all instances in ℬ_*i*_, all the synthetic instances in 𝒮 are merged with the real instances to create a new set of instances 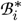 that includes 2*m* instances. From line 13 to line 38, each real instance *x*_*j*_ in ℬ_*i*_ is used as an anchor to generate loss function values. In terms of each anchor *x*_*j*_, five variables 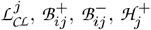 and 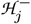 are initialised between lines 14 and 18. Then its corresponding positive and negative instance sets are created by considering the instances in 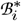 between lines 19 and 28. For each instance *x*_*l*_ other than the anchor, if its class label *y*_*l*_ is different to the class label *y*_*j*_ and the anchor, that instance will be added into the negative sample set 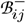 (line 21) and its projection *h*_*j*_ is created (line 22) before being added into 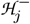 (line 23). *Vice versa*, that instance *x*_*l*_ will be added into the positive instance set of the anchor *x*_*j*_ (line 26). After creating both sets 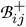 and 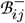, in line 29, all the positive instances in 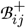 are sorted in ascending order according to the cosine similarity values calculated by *F* (*·*), i.e. the top-ranked instances are the most dissimilar to the anchor. In line 30, a top-*k* percent of the ranked positive instances are used to create another set 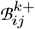, which is used to create the positive instance projection set 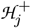 between lines 31 and 34. For each instance *x*_*q*_ in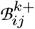, its projection is created by ℰ and 𝒫, and then added into 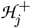. After creating the projection *h*_*j*_ of the anchor in line 35, the loss function value 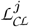 is computed by using 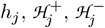 and *τ* (line 36). The value of 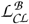 is incremented by the value of 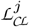 accordingly in line 37. Finally, in line 39, the loss function value 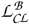 of the batch *B*_*i*_ is normalised by *m* and used to optimize the network parameters of ℰ and 𝒫 (i.e. line 40). The trained encoder network ℰ ^∗^ is returned by Algorithm 1, as shown in line 42.

## 4 Computational Experiments

### 4.1 Experimental Methodology

In this work, we use 18 different scRNA-Seq datasets cover different human and mouse tissues [9], which were collected in [25, 26] to evaluate the performance of GAN-RCL. For each dataset, 20% of instances are used as a held-out validation set that is used for model selection, whilst the remaining 80% of instances are used to conduct GAN-RCL evaluation in a 5-fold cross validation manner. The batch size is set to 64 (i.e. *m*=64), and batch normalisation is adopted for improving training stability. We use a four-layer MLP encoder that consists of three 1024-dimensional hidden layers and a 512-dimensional output layer. The projection head consists of a single 256-dimensional hidden layer and a 128-dimensional output layer. The ReLU activation function is used in both networks, along with the well-known Adam optimizor with a learning rate of 10^−3^ and a weight decay value of 10^−6^. The number of training epochs is set to 500 and the temperature hyperparameter is set to 0.1 (i.e. *τ* = 0.1) as suggested in [11]. On every 5_*th*_ training epoch, we use the frozen encoder to transform the instances in training folds and held-out validation set into feature representations. An SVM classifier is trained using the transformed training folds’ instances to predict the cell-types of the instances in the validation set. Then we select the optimal encoder whose transformed feature representations lead to the highest predictive performance. The grid search approach is also used to select the optimal hyperparameters of the SVM. In terms of the hyperparameter *k* of the proposed GAN-RCL, we select the optimal value that lead the best predictive performance. The value of *k* is drawn from the set {0.1, 0.2, 0.3, 0.4, 0.5, 0.6, 0.7, 0.8, 0.9, 1.0} that covers from selecting the top 10% hardest positive instances up to selecting all positive instances.

In terms of the scGAN training process, we set the batch size as the number of training instances due to the class imbalance issue, this ensures enough instances for all individual classes can be used for the instance selection process. For consistency, we use the same five training folds that are used for the instance selection and contrastive learning processes. The scGAN is composed of a four-layer MLP generator with three 512-dimensional hidden layers. Given the nature of scRNA-seq datasets, the output of the generator is fed to a ReLU function. The discriminator is composed of a four-layer MLP with three 256-dimensional hidden layers. The output of the discriminator is transformed into probabilities using the Sigmoid function. LeakyReLU activation is used in both networks. The training is optimised using Adam with a learning rate of 10^−4^, *β*_1_ = 0.5 and *β*_2_ = 0.9. The class embeddings size is set to 50. The generator training is regularised using the dropout approach whose value is set to 0.2. *γ* is set to 0.3, hence the label smoothing parameter *ϵ* is randomly drawn from the uniform distribution *U* (0, 0.3). The number of training epochs is set to 2,000. After every 40 epochs’ training, we feed the generator with random Gaussian noises along with the class labels of the held-out validation set. This generates synthetic samples that bear the same class labels as the validation sample. Following the work in [16], we conduct classifier-two-sample-tests (CTSTs) to evaluate the performance of the generator network, i.e. the ideal Leave-One-Out Cross-Validation (LOOCV) accuracy of a CTST equals to 0.500, suggesting that the generated synthetic instances come from the same distribution of the training (real) instances. Given the multi-class setting, we conduct CTSTs for each class, then reported the average accuracy across all classes. All code is implemented using PyTorch [27] and Scikit-learn [28]. In terms of the predictive performance evaluation on cell-type identification methods, we use three well-know metrics, i.e. Matthews correlation coefficient value, F1 score and accuracy value, all in their multi-class settings [9], namely MC-MCC, MC-F1 and ACC.

### 4.2 Experimental Results

#### 4.2.1 scGAN successfully learns the scRNA-Seq expression profile distributions of different cell-types

We first report the training quality and the performance of scGAN. In general, the trained scGAN successfully learns the high-dimensional distributions of different scRNA-seq datasets. As shown in Figure 2, the average LOOCV CTST accuracies for almost all 18 datasets over 5-fold cross validation are very close to 0.500, which denotes the perfect scenario, when both the real and the GAN-generated synthetic instances follow the same distribution. For example, only two datasets, i.e. Baron Mouse and Segerstolpe, respectively obtain LOOCV CTST accuracies that are lower than 0.500, i.e. 0.441 and 0.449, respectively. Analogously, only two datasets, i.e. Muraro and Quake 10x Bladder, obtain LOOCV CTST accuracies that are higher than 0.500, i.e. 0.631 and 0.701, respectively.

**Fig. 2.**
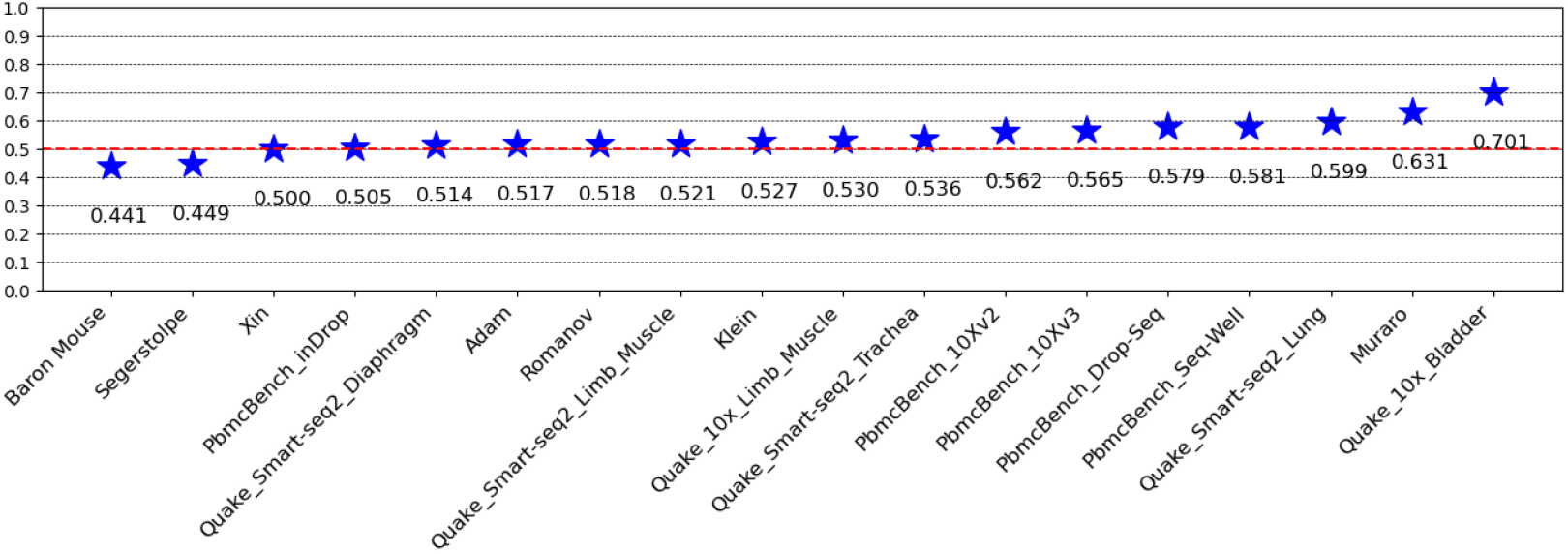
The distributions of leave-one-out cross-validation accuracies of classifier-two-sample-tests averaged over all different classes for 18 individual datasets over 5-fold cross validations.

#### 4.2.2 GAN-RCL successfully improves the predictive performance of AF-RCL and outperforms other state-of-the-art scRNA-Seq contrastive learning-based cell-type identification methods

We then compare GAN-RCL with four different state-of-the-art contrastive learning-based cell-type identification methods, namely AF-RCL-S, AF-RCL [9], Sup-GsRCL [6] and Sup-RGMRCL [6]. AF-RCL-S that is a variant of GAN-RCL also conducts hard positive sample selection but does not exploit any synthetic instances. As shown in Table 2, GAN-RCL, AF-RCL-S and AF-RCL all follow the augmentation-free approach and adopt the modified supervised contrastive learning loss function [9]. Sup-GsRCL and Sup-RGMRCL both use the conventional supervised contrastive learning loss function [11], but adopts different view creation strategies, i.e. adding Gaussian noise and masking random genes, respectively. Table 3 shows the results obtained by those five different methods, whilst Figures 3.A-3.C show the pairwise comparisons between those methods’ performance.

**Table 2.**
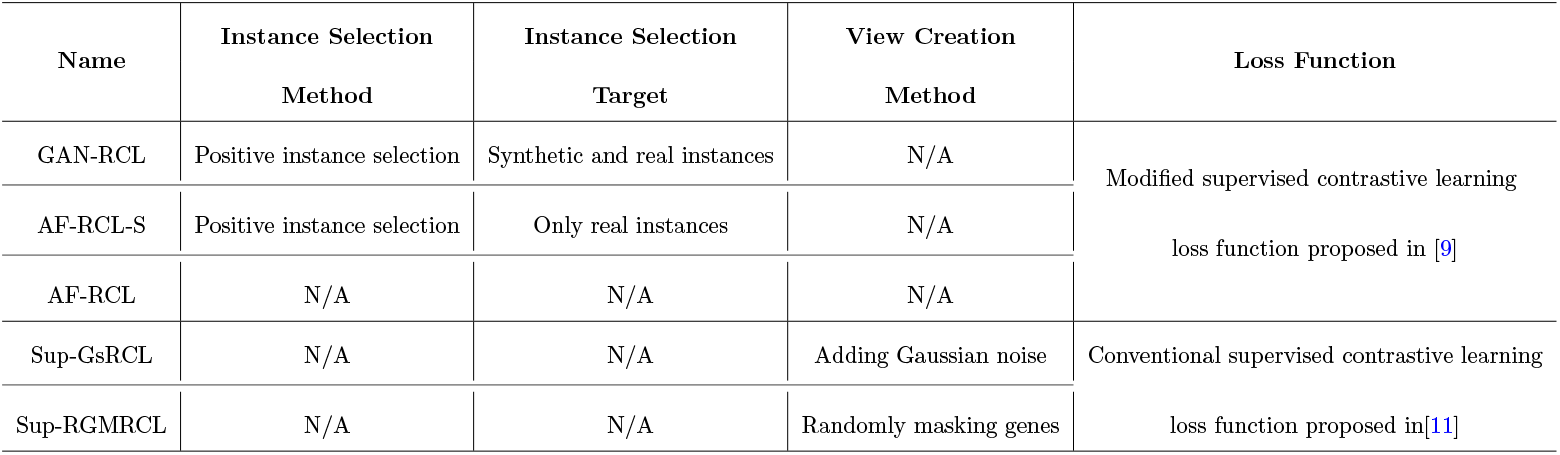
The characteristics of different state-of-the-art supervised contrastive learning-based cell-type identificaiton methods.

**Table 3.**
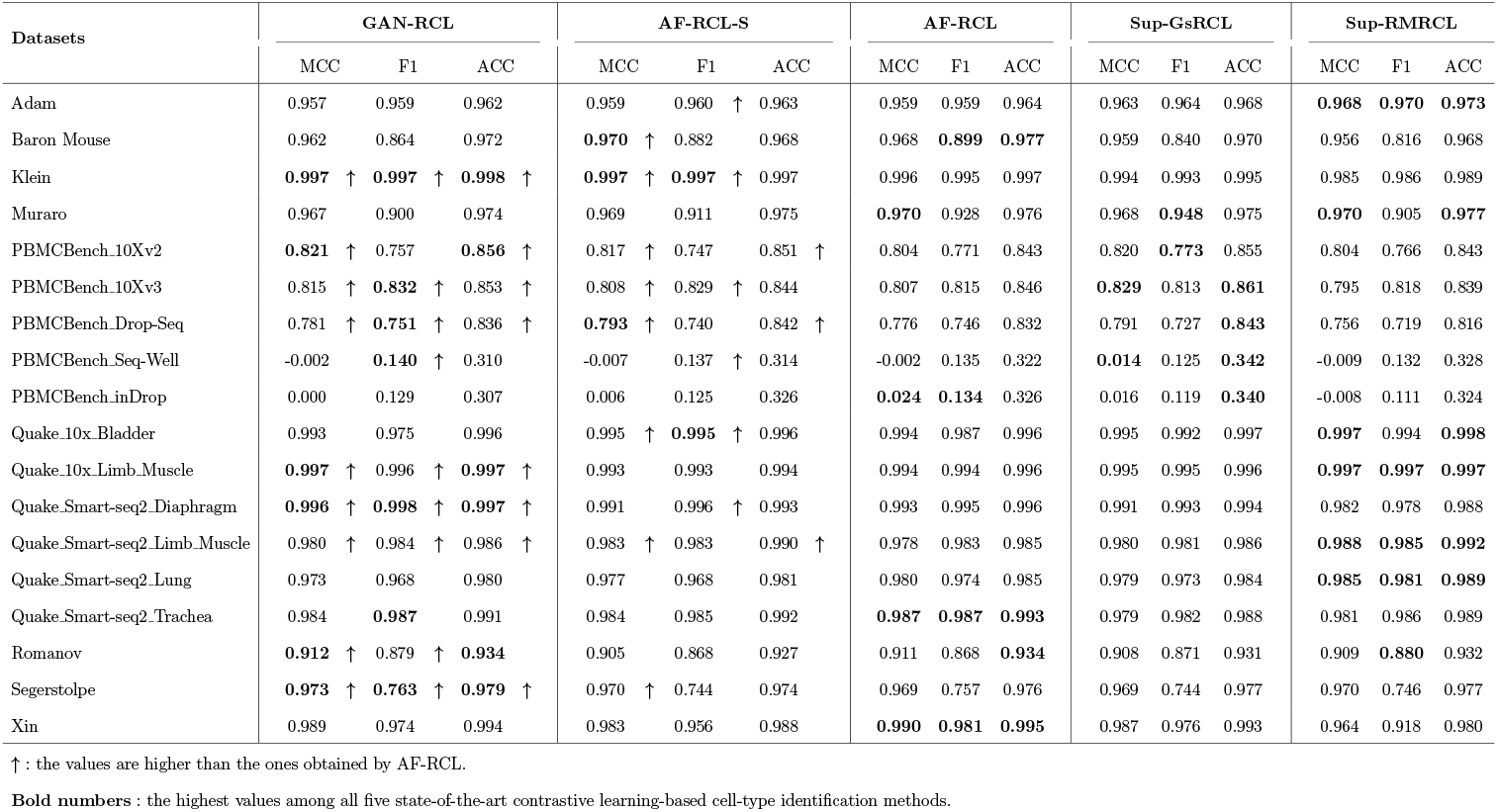
Predictive performance of GAN-RCL and other state-of-the-art contrastive learning-based cell-type identification methods.

**Fig. 3.**
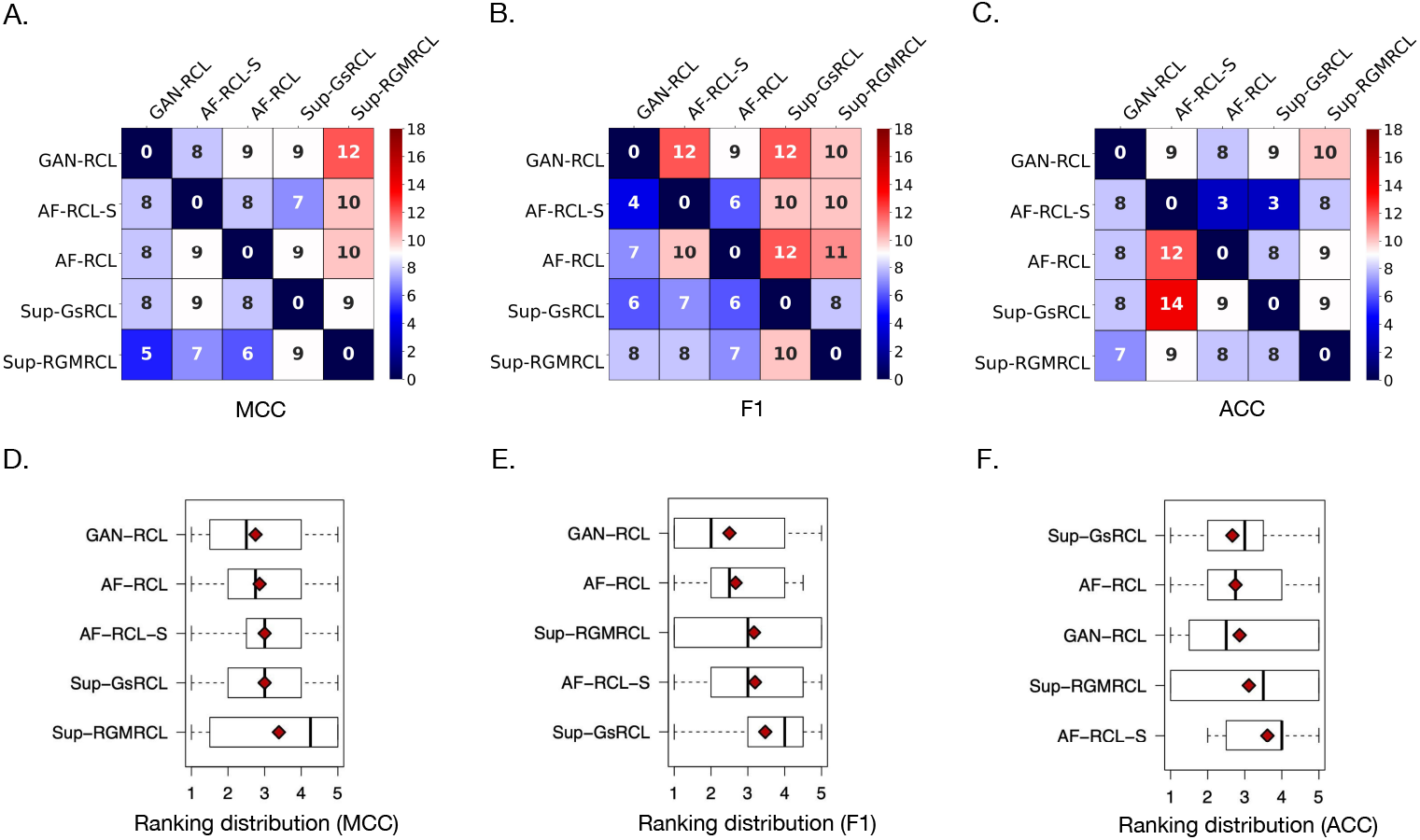
(A–C) The heatmaps showing the pairwise comparisons of five different state-of-the-art contrastive learning-based cell-type identification methods according to the numbers of tasks where each method obtained higher values of the corre-sponding metrics. (D–F) The boxplots showing the ranking distributions of five different state-of-the-art scRNA-Seq supervised contrastive learning-based cell-type identification methods, where the average rankings are denoted by the red diamond symbol.

In general, GAN-RCL successfully improves the performance of AF-RCL and outperforms other supervised scRNA-Seq contrastive learning methods. To begin with, in terms of MCC values and F1 scores, GAN-RCL successfully improves the predictive performance of AF-RCL on 9 out of 18 tasks, as denoted by the arrows in the 2_*nd*_ and the 3_*rd*_ columns of Table 3. It also obtains higher ACC values than AF-RCL on 8 out of 18 tasks, as shown in the 4_*th*_ column. In Figure 3, AF-RCL only performs better than GAN-RCL on 8, 7 and 8 tasks, according to those three different metrics respectively. Both methods obtain the same MCC values, F1 scores and ACC values on 1, 2 and 2 tasks, respectively. Moreover, among those 18 tasks, GAN-RCL outperforms AF-RCL-S on 8, 12 and 9 tasks, according to those three different metrics respectively. In terms of AF-RCL-S, it only performs better than GAN-RCL on 8, 4 and 8 tasks. As a method that conducts hard positive sample selection, it also only improves the predictive performance of AF-RCL on 8, 6, and 3 tasks, according to MCC values, F1 scores and ACC values respectively. Furthermore, GAN-RCL obtains higher MCC and ACC values than Sup-GsRCL on 9 out of 18 tasks. It also outperforms Sup-GsRCL on 12 out of 18 tasks, according to F1 scores. In terms of Sup-GsRCL, it only performs better than GAN-RCL on 8, 6 and 8 tasks, according to those three metrics respectively. Finally, GAN-RCL outperforms Sup-RGMRCL on 12 tasks according to MCC values. It also obtains higher F1 scores and ACC values than Sup-RGMRCL on 10 tasks.

Among all those five state-of-the-art contrastive learning-based cell-type identification methods, GAN-RCL obtains the overall highest MCC values, F1 scores and ACC values on 6, 7 and 6 tasks, whilst Sup-RGMRCL performs the best on 6, 5 and 6 tasks, as denoted by the bold numbers in Table 3. AF-RCL obtains the highest MCC values, F1 scores and ACC values on 4 tasks according all three different metrics, and Sup-GsRCL performs best on 2, 2 and 4 tasks respectively according to MCC values, F1 scores and ACC values. AF-RCL-S only obtains the highest MCC values on 3 tasks and the highest F1 scores on 2 tasks. In addition, Figures 3.D-3.F show the average rankings of different contrastive learning-based cell-type identification methods over those 18 tasks. It is obvious that GAN-RCL obtains the best average rankings according to MCC values and F1 scores, whilst AF-RCL is the second-best performing method, due to its second-best average rankings in all three metrics, though the best ACC-based average ranking is obtained by Sup-GsRCL.

#### 4.2.3 GAN-RCL successfully outperformed other well-known cell-type identification methods

We then compare GAN-RCL with other well-known machine learning-based cell-type identification methods, i.e. scPred [7], SingleCellNet [29] and ACTINN [8]. As show in Figure 4, GAN-RCL outperforms scPred in 9, 10 and 8 tasks according to MCC values, F1 scores and ACC values, respectively. Both methods obtain the same MCC values, F1 scores and ACC values respectively in 2, 1 and 1 tasks. In terms of SingleCellNet, it is outperformed by GAN-RCL in the majority of tasks. GAN-RCL obtains higher MCC values, F1 scores and ACC values respectively in 13, 10 and 14 tasks, whereas SingleCellNet only obtains the same F1 score in 1 task. Analogously, GAN-RCL also outperforms ACTINN in the majority of the 18 tasks. The former obtains higher MCC values, F1 scores and ACC values in 13, 12 and 12 tasks. The latter only obtains the same MCC values, F1 scores and ACC values as the former in 1, 1 and 2 tasks, respectively.

**Fig. 4.**
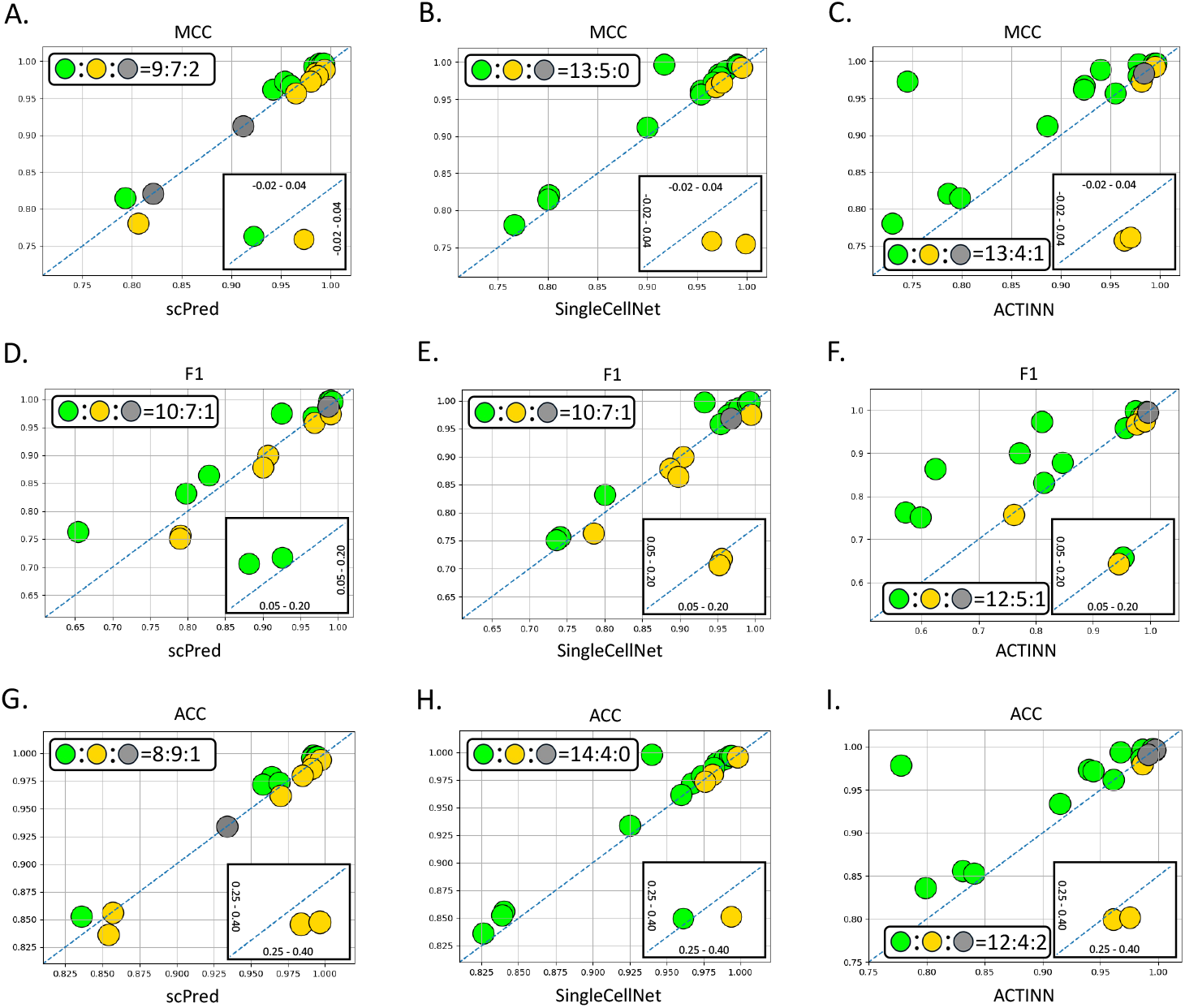
(A–I) The pairwise comparisons between GAN-RCL and other machine learning-based cell-type identification methods.

## 5 Conclusion and Future Research Directions

In this work, we propose a novel generative adversarial networks-enhanced contrastive learning method that conducts hard positive sample selection to improve the quality of hypersphere feature spaces. The proposed GAN-RCL successfully improves the predictive performance of AF-RCL and outperforms other state-of-the-art cell-type identification methods. Future research would be focused on extending the GAN-RCL method on hard negative sample selection tasks to further improve the performance of augmentation-free contrastive learning paradigm on single-cell RNA-Seq analytics.

## Acknowledgement

The authors acknowledge the support of the School of Computing and Mathematical Sciences and the Birkbeck GTA programme.

